# Residual force depression is not related to positive muscle fascicle work during submaximal voluntary dorsiflexion contractions in humans

**DOI:** 10.1101/2023.09.15.557211

**Authors:** Brent James Raiteri, Leon Lauret, Daniel Hahn

## Abstract

Residual force depression (rFD) following active muscle shortening is commonly assumed to strongly and linearly increase with increasing muscle work, but this has not been systematically tested during voluntary contractions in humans. Using dynamometry, we compared steady-state ankle joint torques (*N*=16) following tibialis anterior (TA) muscle-tendon unit (MTU) lengthening and shortening to the torque during submaximal voluntary fixed-end dorsiflexion reference contractions (REF) at a matched MTU length and EMG amplitude. B-mode ultrasound revealed that TA fascicle shortening amplitudes were significantly reduced (*p*<0.001) during MTU lengthening with no preload over small (LEN_small_) and medium (LEN_medium_) amplitudes, respectively, relative to REF. MTU lengthening with a preload over a large (LEN_largeP_) amplitude significantly (*p*<0.001) increased fascicle shortening relative to REF, as well as stretch amplitudes relative to LEN_small_ and LEN_medium_ (*p*≤0.001), but the significant (*p*≤0.028) steady-state fascicle force enhancement relative to REF was similar to LEN_small_ and LEN_medium_ (3-5%). MTU shortening with and without a preload over small (SHO_smallP_/SHO_small_) and large (SHO_largeP/_SHO_large_) amplitudes significantly (*p*<0.001) increased positive fascicle and MTU work relative to REF, but significant (*p*≤0.006) rFD was observed in SHO_smallP_ and SHO_largeP_ (7-10%) only. rFD was linearly related to positive MTU work (*r*_rm_(47)=0.48, *p*<0.001), but not positive fascicle work (*r*_rm_(47)=0.16, *p*=0.277). Our findings indicate that MTU lengthening without substantial fascicle stretch enhances steady-state force output, which might be due to less shortening-induced rFD. Our findings also indicate that different amounts of positive fascicle and MTU work induce similar rFD, which cautions against using work to predict rFD during submaximal voluntary contractions.

## Introduction

Skeletal muscles power a remarkable repertoire of movements in everyday life under submaximal voluntary activation. However, our ability to predict neuromuscular output under these conditions is poor (Dick *et al*., 2017), which directly limits our ability to assess tissue and joint loading and effectively inform injury prevention and rehabilitation strategies, as well as understand motor control (Imani Nejad *et al*., 2020). Inaccurate predictions of neuromuscular output might arise because muscle models typically assume that force output for a given muscle activity level is only dependent on the muscle’s instantaneous length and velocity, whereas growing evidence suggests that the amplitude of muscle shortening under submaximal voluntary activation strongly affects neuromuscular output (Roberts *et al*., 1997), as does the preload force prior to a perturbation (Edman, 1988; Libby *et al*., 2020). Consequently, modified muscle models that account for additional mechanical factors related to the muscle’s previous state or ‘history’ of force production have shown promise in reducing prediction inaccuracies (McGowan *et al*., 2013). Yet, there is a surprising lack of *in vivo* experimental data from submaximal voluntary contractions to inform such models for everyday movement simulations.

A muscle’s steady-state force can accurately be predicted based on the muscle’s active length alone during fixed-end supramaximal tetanic contractions (Gordon *et al*., 1966). But the same is not true when a dynamic phase precedes the fixed-end hold phase of a contraction. Accurately predicting force during dynamic-hold conditions is challenging because a variable residual force enhancement (rFE) or residual force depression (rFD) exists following active muscle lengthening or shortening (Abbott & Aubert, 1952), respectively, and the magnitude of rFE or rFD depends on different mechanical factors. rFE seems to primarily depend on the muscle’s relative length following active lengthening as a percentage of its optimum length for isometric force production, with *ex vivo* and *in vivo* rFE both increasing with increasing muscle length (Schachar *et al*., 2004; Hisey *et al*., 2009; Bakenecker *et al*., 2020, 2022). Conversely, rFD seems to primarily depend on the preceding muscle work during shortening, with *ex vivo* and *in vivo* rFD increasing with increasing positive muscle or joint work, respectively (Granzier & Pollack, 1989; Herzog *et al*., 2000; Dargeviciute *et al*., 2013).

Interestingly, a positive rFD-work correlation does not exist following stretch-shortening cycles (Seiberl *et al*., 2015; Fortuna *et al*., 2017, 2018, 2019; Hahn & Riedel, 2018; Tomalka *et al*., 2020; Groeber *et al*., 2020, 2021; Fukutani & Herzog, 2021), and worryingly, the reported strong and positive rFD-work correlations were not based on direct measurements of muscle shortening, except in one study (Granzier & Pollack, 1989). Additionally, muscle work was quantified during the externally-imposed shortening only, and not during active force development before the perturbation nor during the fixed-end reference contractions. Some rFD-work analyses also appear to include non-independent observations and aggregated data, which violates the assumption of independence (Bakdash & Marusich, 2017), and could bias the correlations.

The positive rFD-work correlation that is commonly assumed is therefore potentially flawed and raises questions about whether such a relation should be included within modified muscle models designed to improve force prediction accuracy during everyday movement simulations. For example, during maximal voluntary contractions, rFD was independent of the preceding shortening amplitude and speed (Lee *et al*., 2000), whereas rFD should increase with increasing shortening amplitude and increasing mean force (i.e. decreasing shortening speed) if it is best predicted by work. rFD also only increased with increasing shortening amplitude following some, but not all amplitudes, during submaximal voluntary contractions of the human adductor pollicis (Rousanoglou *et al*., 2007). While rFD following submaximal voluntary stretch-shortening cycles (SSCs) has yet to be investigated, studies using artificial muscle activation found no positive correlation between rFD and net joint work because rFD was reduced despite higher net joint work during shortening of SSCs relative to shortening-hold conditions without prior stretch (Seiberl *et al*., 2015; Fortuna *et al*., 2017, 2018, 2019; Hahn & Riedel, 2018; Groeber *et al*., 2020, 2021). These findings thus raise questions about whether there is a generalisable positive rFD-work correlation across different contraction conditions.

Contraction conditions that do not typically induce rFD involve small shortening amplitudes and no preload before shortening (Edman, 1966; Gordon *et al*., 1966; Edman *et al*., 1993). While it could be argued that rFD was minimal because positive muscle work was low in these conditions (Granzier & Pollack, 1989), another potential and typically overlooked reason for negligible rFD is that the fixed-end reference (REF) contractions used to estimate rFD are contaminated with rFD themselves. We previously showed that by reducing the amount of active muscle shortening via muscle-tendon unit (MTU) lengthening of the active muscles during a torque ramp, steady-state active torque was enhanced relative to that in the REF condition at a similar muscle activity levels and fascicle lengths (Raiteri & Hahn, 2019). Consequently, we concluded that the active muscle shortening permitted by effectively greater in-series compliance in the REF condition compared with the MTU lengthening-hold condition could induce rFD.

More recent findings from the *in situ* rabbit tibialis anterior muscle support our results (Mahmood *et al*., 2021), but those authors interpreted their findings differently. Mahmood and colleagues (2021) suggested that the enhanced steady-state active force following MTU lengthening with simultaneous muscle fibre shortening was due to mechanisms underlying rFE rather than rFD. This interpretation was justified based on an assumed positive rFD-muscle work correlation, and their finding of similar positive muscle work production between the MTU lengthening-hold and REF conditions. However, as stated above, positive muscle work does not always predict rFD accurately. Additionally, it is hard to envision how the typically observed active and passive contributions to rFE would enhance the steady-state force following active fibre shortening because rFE is typically observed following active muscle fibre stretch and rFE increases with increasing muscle lengths beyond the optimum length for isometric force production (Herzog & Leonard, 2002, 2005; Schachar *et al*., 2002, 2004). Consequently, we believe it is more likely that reduced rFD contributes to the enhanced steady-state force following simultaneous MTU lengthening and fibre shortening.

We designed the following human research study with three aims in mind. We wanted to: 1) determine whether shortening during the initial force rise of fixed-end contractions (REF) of a moderately-compliant MTU induces rFD, 2) investigate how preload affects rFD following MTU shortening-hold conditions compared with REF, and 3) examine the relation between rFD and positive muscle (i.e. fascicle) and MTU work. To test our aims, we opted for submaximal voluntary contractions because they are more physiologically relevant for everyday movement than maximal voluntary and non-voluntary conditions. Further, we studied the human tibialis anterior (TA) muscle specifically because: 1) a joint-angle-specific muscle-tendon moment arm function is available from the literature (Maganaris *et al*., 1999); 2) TA contributes 45-52% of the maximal voluntary dorsiflexion torque as assessed via electrical stimulation of its muscle belly (Maganaris & Paul, 2000; De Zee & Voigt, 2002), and; 3) the entire length of TA’s muscle fascicles are visible within a 4-8 cm ultrasound image field of view (Ito *et al*., 1998; Maganaris *et al*., 2001). We imposed MTU lengthening and MTU shortening with and without a preload over different amplitudes to systematically vary the amount of TA fascicle work before a steady-state hold phase was reached at a matched MTU length and TA muscle activity level between conditions.

We hypothesised that: 1) steady-state active torques following MTU lengthening would be enhanced relative to REF because of reduced muscle fascicle shortening amplitudes during the torque ramp, which was previously observed (Raiteri & Hahn, 2019; Mahmood *et al*., 2021); 2) steady-state active torques following MTU shortening with and without a preload would be and would not be depressed relative to REF, respectively, as previously shown (Edman *et al*., 1993), and; 3) the estimated steady-state active force differences following MTU shortening normalised to REF (i.e. rFD) would be strongly and positively correlated with the MTU work during shortening, in line with previous findings (Granzier & Pollack, 1989; Herzog *et al*., 2000), but rFD would not be positively correlated with muscle fascicle work during shortening.

## Methods

### Ethical approval

The experimental protocol and procedures were approved by the Ethics Committee of the Faculty of Sport Science at Ruhr University Bochum (EKS V 33/2019). The study conformed to the standards set by the latest version of the Declaration of Helsinki, except for database registration. Following verbal and written explanation of all procedures and risks, free written informed consent was given by each participant prior to participating in the study.

### Sample size calculation

Simulation-based power analysis was performed in RStudio (v1.4.1717, 2021, Boston, Massachusetts, USA) using Superpower (Lakens and Caldwell, 2021) to determine the minimum sample size required to achieve at least 80% power to detect a minimum effect size of interest (*d*_z_) of 0.6 between estimated marginal means with a family-wise two-tailed alpha level of 5%. Results from Table 1 of Tilp et al. (2011) were simulated as it is the only *in vivo* study we are aware of that investigated steady-state dorsiflexion torques following MTU lengthening and shortening of different magnitudes and velocities. The ANOVA design was set to one within-subject factor with eight levels, and a multivariate model was used with a Holm-Bonferroni correction for estimated marginal mean comparisons relative to the REF condition. The means for each condition were based on reported values of: REF = 0; small-magnitude MTU lengthening or LEN_small_ = 7 (i.e. minimum rFE), medium-magnitude MTU lengthening or LEN_medium_ = 10 (i.e. mean rFE), large-magnitude preloaded MTU lengthening or LEN_largeP_ = 13 (i.e. maximum rFE); small-magnitude MTU shortening or SHO_small_ = 0; small-magnitude preloaded MTU shortening or SHO_smallP_ = −11 (i.e. minimum rFD), large-magnitude MTU shortening or SHO_large_ = 0; large-magnitude preloaded MTU shortening or SHO_largeP_ = −21 (i.e. maximum rFD). A common standard deviation of 11.67 was expected, which represents the mean standard deviation of the rFE and rFD results from Tilp et al. (2011), and a correlation of at least 0.82 between repeated measurements was predicted.

From 10,000 simulations, a sample size of 14 was estimated to have 100% power to detect a Cohen’s *f* of 0.85, and over 82% power to detect a minimum Cohen’s *d*_z_ of 1.11 between LEN_small_ and REF with a family-wise alpha level of 5%. A sample size of 14 also resulted in over 98% power to detect a minimum Cohen’s *d*_z_ of 1.58 between the other comparisons that were expected to be different; a seed number of 2022 can be used to replicate these results. As reducing the sample size to 13 resulted in less than 79% power to detect a significant effect between LEN_small_ and REF following a Holm-Bonferroni correction for multiple comparisons, a sample size of 14 was deemed as the minimum necessary to sufficiently power this study.

### Participants

Sixteen healthy participants (eight women, age: 26.3±2.6 years; height: 177.3±9.6 cm; mass: 72.2±12.8 kg) without recent lower limb injury or surgery were recruited for this study. Participants were physically active at the time of the study and had not previously systematically trained their dorsiflexor muscles.

### Experimental setup

The experimental setup is illustrated in Figure 1 of Raiteri et al. (2023) and will only be briefly described here. Participants sat in a reclined posture (∼115° backrest relative to seat) while they performed voluntary contractions with their right dorsiflexors on a motorized dynamometer. A dorsiflexion attachment over the metatarsals ensured that the plantar aspect of their foot maintained constant contact with the dorsi/plantar flexion adapter of the dynamometer and was also used to limit ankle joint rotation and force contributions from the toe extensors to the measured net ankle joint torque. The right foot and thigh were vertically aligned in the frontal plane, and the right knee and hip were at approximately 90° and 110° of flexion in the sagittal plane, respectively.

**Figure 1.**
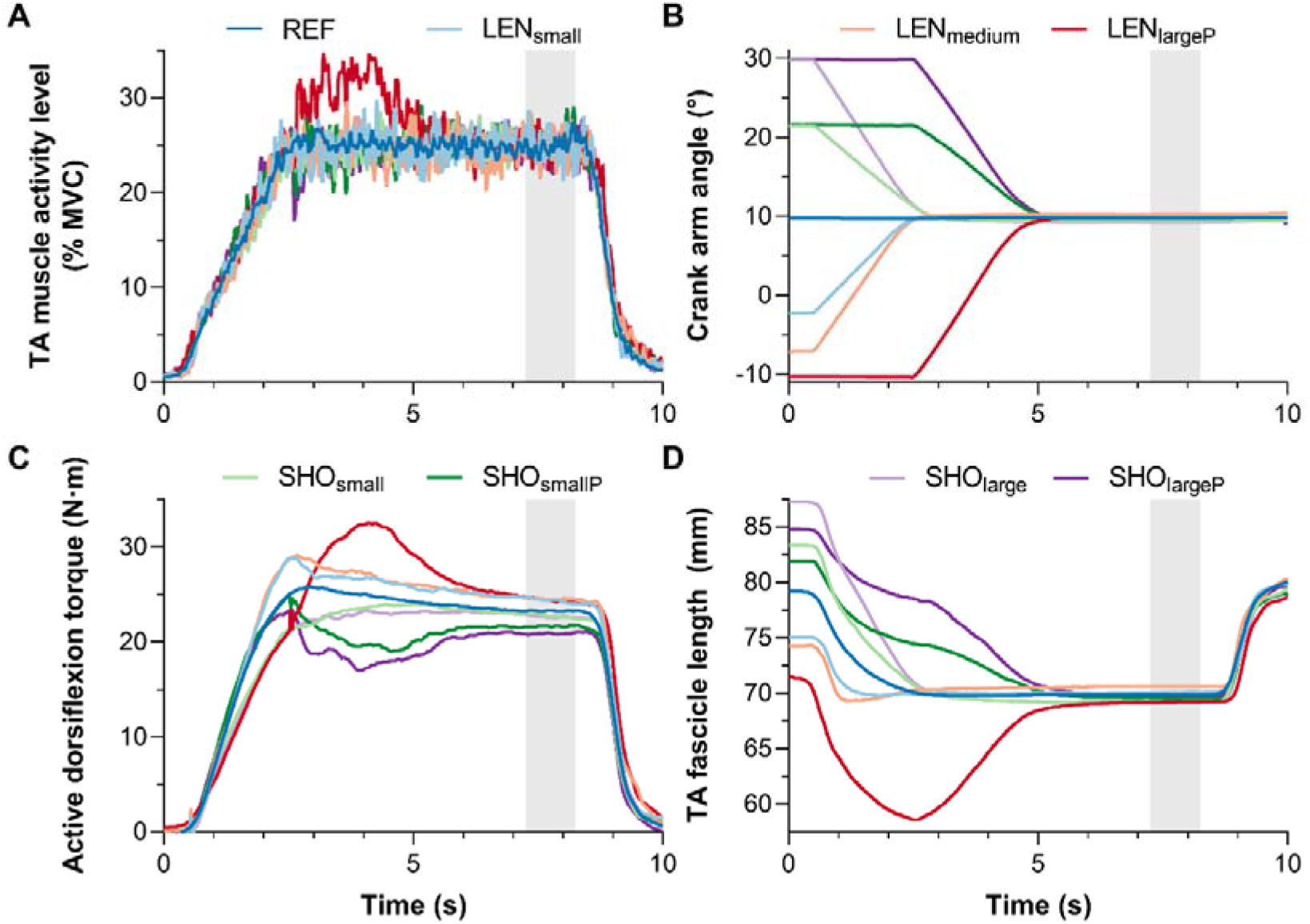
Mean (A) normalised tibialis anterior (TA) muscle activity level-time, (B) crank arm angle-time, (C) active dorsiflexion torque-time, and (D) TA muscle fascicle length-time traces from sixteen participants during fixed-end reference (REF), and muscle-tendon unit (MTU) lengthening (LEN)-hold and shortening (SHO)-hold conditions with and without a preload (_P_) over small, medium, and large amplitudes in EMG-amplitude-matched contractions. Different steady-state active torques were clearly produced (C) at similar muscle activity levels (A), muscle-tendon unit lengths (B), and fascicle lengths (D). The grey shaded areas indicate the analysed one-second steady-state phase of the contractions. Note the relatively negligible muscle fascicle stretch (C) in LEN_small_ during the imposed MTU lengthening (B). Also note that plantar flexion angles are positive and there was 1 missing dataset for 2 separate participants in LEN_small_ and LEN_medium_.

The experimental techniques are identical to those described in Raiteri et al. (2023) and are described in detail there. Briefly, a motorized dynamometer (IsoMed2000, D&R Ferstl GmbH, Hemau, Germany) measured net ankle joint torque and the angle of the footplate relative to the fixed right shank at 2 kHz using a 16-bit analogue-to-digital converter (Power1401-3) and Spike2 (8.23 64-bit version) data collection system (Cambridge Electronic Design Ltd., Cambridge, United Kingdom). The data collection system used a ±5 V input range and recorded all digital signals synchronously. A flat linear-array ultrasound transducer (LV8-5N60-A2, TELEMED, Vilnius, Lithuania) was attached over TA’s mid-belly with a constant pressure using self-adhesive bandage (7.5 cm width, COJJ, Amazon, Seattle, Washington, United States) to image TA’s superficial and deep compartments with a 60 (width) × 50 (depth) mm field of view at ∼34 fps using a coupled PC-based ultrasound system (ArtUs EXT-1H, TELEMED). Surface electrodes (hydrogel Ag/AgCl, 8 mm recording diameter, H124SG, Kendall International Inc., Mansfield, Massachusetts, United States) were secured to the skin (1.25 cm width surgical tape, 3M Transpore, St. Paul, Minnesota, United States) above TA’s distal superficial compartment in a bipolar configuration (2 cm inter-electrode distance) and above the right fibular head following standard skin preparation. These electrodes were used to record TA’s single differential myoelectric signal at 2 kHz using surface electromyography (NL905, Digitimer Ltd., Welwyn Garden City, United Kingdom).

### Experimental protocol

Each participant completed one experimental session. Participants first preconditioned their dorsiflexor MTUs by performing five submaximal (1 s hold, 1 s rest, ∼80% of perceived maximum) voluntary dorsiflexion contractions at 15° plantar flexion (Maganaris *et al*., 2002). After two minutes of rest, participants then performed one to five maximal (3-s to 5-s hold) voluntary dorsiflexion contractions at a minimum of five plantar flexion angles ranging from 0-30° plantarflexion (where 0° plantar flexion = footplate perpendicular to shank) to determine their angle of maximum dorsiflexion torque production. Participants received real-time visual feedback of their net ankle joint torque and standardised verbal encouragement from the investigator (e.g., pull the top of your foot towards your shank as hard as possible using only your dorsiflexors). At least two minutes of rest was provided to minimise fatigue following each maximal voluntary contraction.

After five minutes of rest, participants completed eight TA EMG-amplitude-matched submaximal voluntary conditions (Fig. 1A&B) that consisted of one fixed-end REF condition, three MTU lengthening-hold conditions, and four MTU shortening-hold conditions. The TA EMG signal was displayed on a screen in front of participants as a real-time (125 ms delay) and smoothed signal (i.e. the DC offset was removed and the moving 250 ms moving root-mean-square amplitude was calculated) so that participants could attempt to visually match their TA muscle activity level to within predefined traces that were 10% of the maximum EMG amplitude apart. After a 2-s ramp phase, these predefined traces were fixed at 20% and 30% of the maximum EMG amplitude recorded at the reference MTU length; the visual gain of the EMG feedback was maintained throughout the experiment.

The reference MTU length, which was common to all eight conditions, was individualised as the angle of maximum dorsiflexion torque production. The reference length was individualised to reduce variability in the rFD results because rFD appears to be muscle-length dependent (Morgan *et al*., 2000; Van Noten & Van Leemputte, 2011), and this reference length was chosen to maximise the capacity of the dorsiflexors to produce work under submaximal voluntary conditions. The REF condition involved a 2-s ramp and then a 6-s hold phase at the desired muscle activity level and reference MTU length, which allowed a steady-state torque to be achieved. The REF condition was performed after two consecutive trials from each of the seven dynamic conditions.

The seven dynamic conditions had a variable crank arm rotation velocity to ensure that the duration of the dynamic phase was fixed at 2 s. The first and second dynamic conditions involved MTU lengthening without a preload over a small (11-14°; LEN_small_) or medium amplitude (15-20°; LEN_medium_) during a 2-s ramp, and the ramp was followed by a 6-s hold phase at the desired muscle activity level and reference MTU length. The 11° amplitude was the smallest rotation that the dynamometer could perform, but despite setting equivalent parameters in the software, the rotations were not always identical (hence the 3° range). LEN_small_ was always performed prior to LEN_medium_; with the aim to prevent fascicle stretch during the rotation. LEN_medium_ generally reduced the amount of fascicle shortening more than LEN_small_, but with an increased likelihood of fascicle stretch during the rotation. The third dynamic condition involved a fixed-end preload (_P_) phase, which was developed during a 2-s ramp, before MTU lengthening over a large amplitude (19-22°; LEN_largeP_), and then a 4-s hold phase at the desired muscle activity level and reference MTU length. The large amplitude (i.e. mean of 21°) was designed to be 10° larger than the small amplitude, but no large-amplitude MTU lengthening without a preload was performed as this typically resulted in unwanted fascicle stretch during the rotation.

The last four dynamic conditions were randomized and involved MTU shortening without a preload (i.e. 2-s ramp, 6-s hold) and MTU shortening with a preload (i.e. 2-s ramp, 2-s rotation, 4-s hold) over a small (11-13°; SHO_small_ and SHO_smallP_) or large (19-22°; SHO_large_ and SHO_largeP_) amplitude. The MTU shortening-hold conditions were designed to mirror the MTU lengthening-hold conditions, with the exception that no medium-amplitude MTU shortening-hold conditions were performed to reduce the risk of fatigue. A trial from a dynamic condition was considered invalid and repeated if the muscle activity level was not within the predefined traces (i.e. ±5% of the desired 25% muscle activity level) for the duration of the contraction, and trials were repeated until there were at least two valid contractions from each dynamic condition. At least 90 s of rest was provided to minimise fatigue after each submaximal voluntary contraction. Following the submaximal voluntary contractions, a further two minutes of rest was provided, and six passive ankle rotations were performed at 5°·s^-1^ over each participant’s ankle joint range of motion.

### Data processing and analysis

The first data processing step involved using a custom-written script (spike2mat.s2s; doi:10.5281/zenodo.7411280) in Spike2 to export individual trials from a recording of the entire experimental session as separate files with a .mat extension. The recorded digital signals from each trial were cropped between a common start and end time using the timestamps of the first and last digital pulses from each ultrasound video recording. The cropped digital signals from each trial were then combined with the tracked fascicle data from the corresponding ultrasound video recording; data was considered synchronized only when the duration difference between ultrasound and digital files was no more than one frame. The tracked fascicle data included absolute lengths and angles (relative to the horizontal) of representative fascicles from TA’s superficial and deep muscle compartments from each ultrasound image, which were determined in MATLAB (R2022a 64-bit version, MathWorks, Natick, Massachusetts, United States) using an updated version (UltraTrack_v5_3.m; doi:10.5281/zenodo.7411280) of UltraTrack (Farris and Lichtwark, 2016). Further details about the absolute fascicle length and angle determinations can be found in Text S1.

Once the cropped digital signals and tracked fascicle data were within the same file, a time vector was constructed using the timestamps of the torque signal and each signal was subsequently resampled using linear interpolation. Torque and angle data were filtered using zero-lag second-order 20 Hz and 6 Hz low-pass Butterworth filters, respectively, which were corrected for two passes (Winter, 2009). To calculate active torque, a steady-state passive net ankle joint torque-angle fit was constructed, then evaluated at the crank arm angles recorded during each trial and subtracted from the recorded net ankle joint torque. Further details about the active torque calculation can be found in Text S2. TA’s muscle activity level was calculated by rectifying the recorded EMG signal after the DC bias was removed and the rectified signal was then smoothed with the same Butterworth filter described above, but at a 10 Hz low-pass cut-off frequency.

From each trial of interest, the steady-state phase was defined as a 1-s period from 9.75 s after the start of the ramp. The means of all outcome variables of interest (e.g., muscle activity level, active torque, TA fascicle length and positive as well as net fascicle work) were calculated over this steady-state period for each valid (i.e. EMG-amplitude matched) trial, and then the mean of the valid steady-state means was calculated for each condition. For the REF condition, if there was a torque difference ≥ 1.0 Nm between the steady state and the previous 1-s period (i.e. from 8.75 s), then the trial was excluded from the analysis.

Active dorsiflexor force (F_active_) was estimated by:

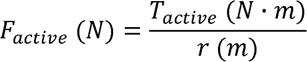

where *T*_*active*_ represents active torque, and *r* represents TA’s joint-angle-specific, MRI-based muscle-tendon moment arm length measured at rest (Maganaris *et al*., 1999). TA tendon force was then estimated by assuming a fixed 50% contribution of TA to the total dorsiflexor force (Brand *et al*., 1986; Maganaris & Paul, 2000; De Zee & Voigt, 2002):

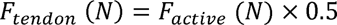

and TA muscle fascicle force was estimated by:

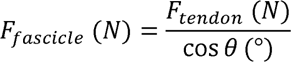

where *0* represents the respective fascicle angle from TA’s superficial compartment. The steady-state normalised active force difference (*F*_*diff*_) was calculated by:

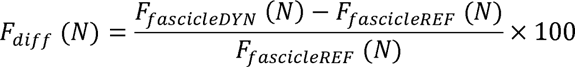

where *F*_*fascicleDYN*_ refers to the mean estimated steady-state active force from one of the dynamic conditions and *F*_*fascicleREF*_ refers to the mean from the REF condition. The same calculation was used to determine the steady-state normalised active torque difference.

TA MTU and fascicle work magnitudes were estimated using the trapezoidal method as the area under the MTU force-MTU displacement and fascicle force-fascicle displacement curves, respectively. The curves started 0.25 s before the start of the ramp (i.e. when active torque/force and MTU/fascicle displacement were zero) and ended at the start of the steady state (i.e. 9.75 s). Net work was partitioned into positive and negative work by calculating the areas under the curves when displacement was positive (i.e. MTU/fascicle shortening) or negative (i.e. MTU/fascicle lengthening), respectively. TA fascicle shortening amplitude was calculated as the difference between fascicles lengths at the instant of fascicle shortening and the minimum length attained during contraction. The instant of fascicle shortening was determined as the last zero crossing of the fascicle velocity vector before maximum shortening speed was achieved.

Fascicle stretch amplitudes during the crank arm rotation were calculated in two ways: 1) as the maximum local difference between fascicle lengths while fascicle velocity remained negative, and 2) as the maximum net difference between fascicle lengths after the minimum length was attained and the following maximum length. The larger of these two values was taken as the fascicle stretch amplitude. Only trials with the smallest stretch amplitudes from LEN_small_ and LEN_medium_ were further analysed. If the same ‘smallest’ stretch amplitude occurred in multiple trials, then these trials were averaged unless there was clear passive fascicle lengthening at the start of the rotation because the participant incorrectly timed the activation of their TA. Additionally, only trials with stretch amplitudes <1 mm from the MTU shortening-hold conditions were analysed.

### Statistics

Statistical analysis was performed with GraphPad Prism (9.1.2 64-bit version, San Diego, California, USA) and the alpha level was set at 5%. One repeated-measures mixed-effects analyses with the Greenhouse-Geisser correction were performed to identify mean differences in steady-state muscle activity level, active torque, the active force percent difference relative to REF, and fascicle length. The same tests were performed to identify mean differences relative to REF in fascicle shortening and stretch amplitudes, as well as net fascicle and MTU work production between conditions. Following a significant effect, individual-variance-based Holm-Sidak multiple comparisons were performed between each dynamic condition and REF. Paired *t*-tests were also performed to identify mean differences in the normalised muscle activity level and normalised active torque between the first and last valid REF trials, which would indicate fatigue or learning effects over the course of the experiment. Repeated-measures Pearson correlation coefficients were calculated to test the strength of the relations between the steady-state percent active force difference relative to REF and 1) fascicle stretch amplitude (MTU lengthening-hold conditions only), 2) net fascicle and MTU work production (MTU shortening-hold conditions only) using the rmcorr package (Bakdash & Marusich, 2017). Data are presented as mean ± standard deviation below unless stated otherwise.

## Results

### Exclusions

Ten trials (range: 8-12) from the REF condition were performed and eight trials were analysed on average because trials that exhibited a torque difference ≥1.0 Nm between the steady state and the previous 1-s period were excluded. Although six trials (range: 2-9) from LEN_small_ and LEN_medium_ were performed on average, only one trial (range: 0-2) was analysed on average, which had the smallest fascicle stretch amplitude during the rotation. No valid trials were analysed from LEN_small_ (P7) or LEN_medium_ (P11) for two separate participants. Two trials were performed and analysed on average from the remaining dynamic conditions (i.e. LEN_largeP_, SHO_small_, SHO_smallP_, SHO_large_, SHO_largeP_). Unless otherwise indicated, the results below are based on *n* = 16.

### Fatigue

The mean steady-state muscle activity level was not significantly different between the first and last valid REF trials (mean difference = 0.4±2.1% of the maximum angle-specific activity level, *t*_15_ = 0.72, *p* = 0.485), and neither was the mean steady-state active torque (mean difference = 1.5±3.2% of the maximum angle-specific torque, *t*_15_ = 1.84, *p* = 0.085).

### Muscle activity level

The mean steady-state muscle activity level was not significantly different between conditions, which indicates that the contractions were EMG-amplitude matched (mean differences = ≤0.6% of the maximum angle-specific activity level, *F*_3.66,53.77_ = 0.64, *p* = 0.621, 2 missing values). The between-subject mean muscle activity levels over time from each of the eight conditions are shown in Figure 1A.

### Active torque and fascicle force

The mean steady-state active torque was significantly different between conditions (*F*_2.39,35.10_ = 16.07, *p* < 0.001), which indicates that contraction history affected the dorsiflexors’ torque-producing capacity, despite similar muscle activity levels and crank-arm angles. Active torque was significantly higher by ∼4% relative to REF in LEN_small_ (*d*_z_ = 0.77, *p* = 0.033, *n* = 15), LEN_medium_ (*d*_z_ = 1.43, *p* < 0.001, *n* = 15), and LEN_largeP_ (*d*_z_ = 1.05, *p* = 0.004), and active torque was significantly lower relative to REF by ∼9% in SHO_smallP_ (*d*_z_ = 0.97, *p* = 0.007) and SHO_largeP_ (*d*_z_ = 1.35, *p* = 0.001; Fig. 2). However, active torque was different by only ∼2% relative to REF in SHO_small_ (*d*_z_ = 0.32, *p* = 0.230) and SHO_large_ (*d*_z_ = 0.47, *p* = 0.181; Fig. 2), and these mean differences were not significant. The between-subject mean active torques over time from every condition are shown in Figure 1C.

**Figure 2.**
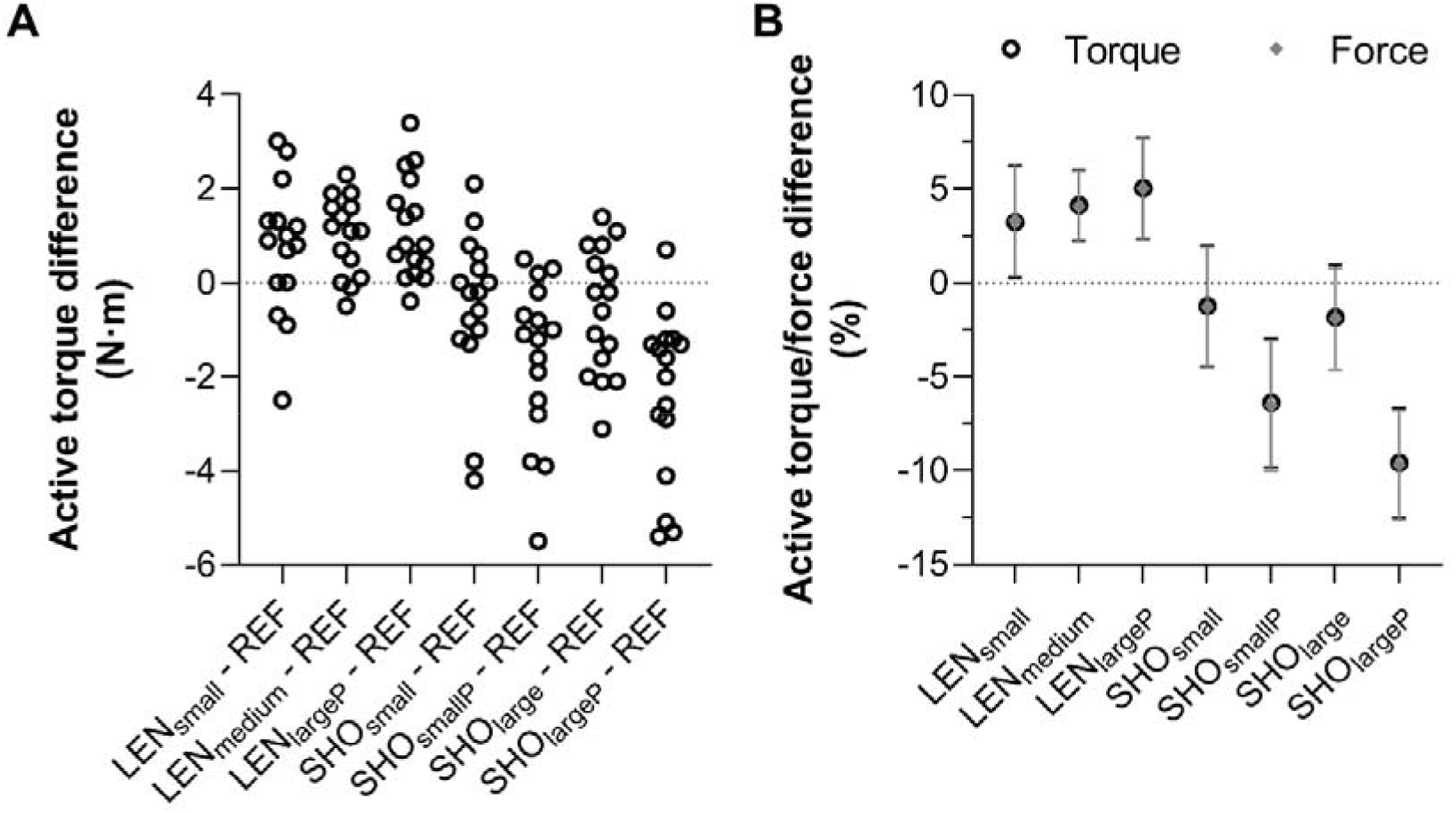
(A) Individual absolute and (B) mean (± 95% confidence intervals) percent differences in steady-state active (A&B) dorsiflexion torque (black open circles) and (B) tibialis anterior force (grey filled diamonds) normalised to the fixed-end reference (REF) condition (horizontal dotted line) following muscle-tendon unit lengthening (LEN) and shortening (SHO) with and without a preload (_P_) over small, medium, and large amplitudes. The 95% confidence intervals show that the active torque and force percent differences relative to REF were not significant for SHO_small_ and SHO_large_ only.

Similar to active torque, the mean steady-state active force was significantly different between conditions (*F*_2.42,35.64_ = 16.01, *p* < 0.001), and the mean steady-state active torque percent differences relative to REF were similar to the mean steady-state active force percent differences (Fig. 2B). The higher mean active forces following MTU lengthening were not significantly different between LEN_small_ (3.3±5.4%, range: −6.7 to 10.4%, *n* = 15, *p* ≥ 0.892), LEN_medium_ (4.1±3.4%, range: −2.6 to 8.6%, *n* = 15), and LEN_largeP_ (5.0±5.1%, range: −2.5 to 14.7%), and the lower mean active forces following preloaded MTU shortening were not significantly different between SHO_smallP_ (−6.5±6.5%, range: −21.5 to 2.1%, *p* = 0.561) and SHO_largeP_ (−9.7±5.5%, range: −18.3 to 2.9%). The mean active forces following MTU shortening without a preload were also not significantly different between SHO_small_ (−1.3±6.0%, range: −12.8 to 12.9%, *p* = 0.922) and SHO_large_ (−1.9±5.1%, range: −9.2 to 6.1%).

### Fascicle length

The mean steady-state fascicle length was not significantly different between conditions (mean differences = ≤1 mm, *F*_1.53,22.58_ = 1.53, *p* = 0.237, 2 missing values), which indicates that differences in fascicle length cannot primarily explain the differences in active torque and force between conditions. The between-subject mean fascicle lengths from every condition over time are shown in Figure 1D.

### Fascicle stretch and shortening amplitudes

The mean fascicle stretch amplitude was significantly different between MTU lengthening-hold conditions (*F*_1.08,15.18_ = 157.2, *p* < 0.001). The mean stretch amplitude was largest in LEN_largeP_ (10.0±3.1 mm, range: 4.2-15.3 mm, *p* ≤ 0.001), followed by LEN_medium_ (1.3±1.0 mm, range: 0.2-3.4 mm, *n* = 15), and LEN_small_ (0.5±0.4 mm, range: 0.1-1.4 mm, *n* = 15; Fig. 3). No significant repeated-measures linear relation was observed between the steady-state active force percent differences relative to REF and fascicle stretch amplitudes between MTU lengthening-hold conditions (*r*_rm_(29) = 0.20, 95% CI = −0.17 to 0.52, unadjusted *p* = 0.288).

**Figure 3.**
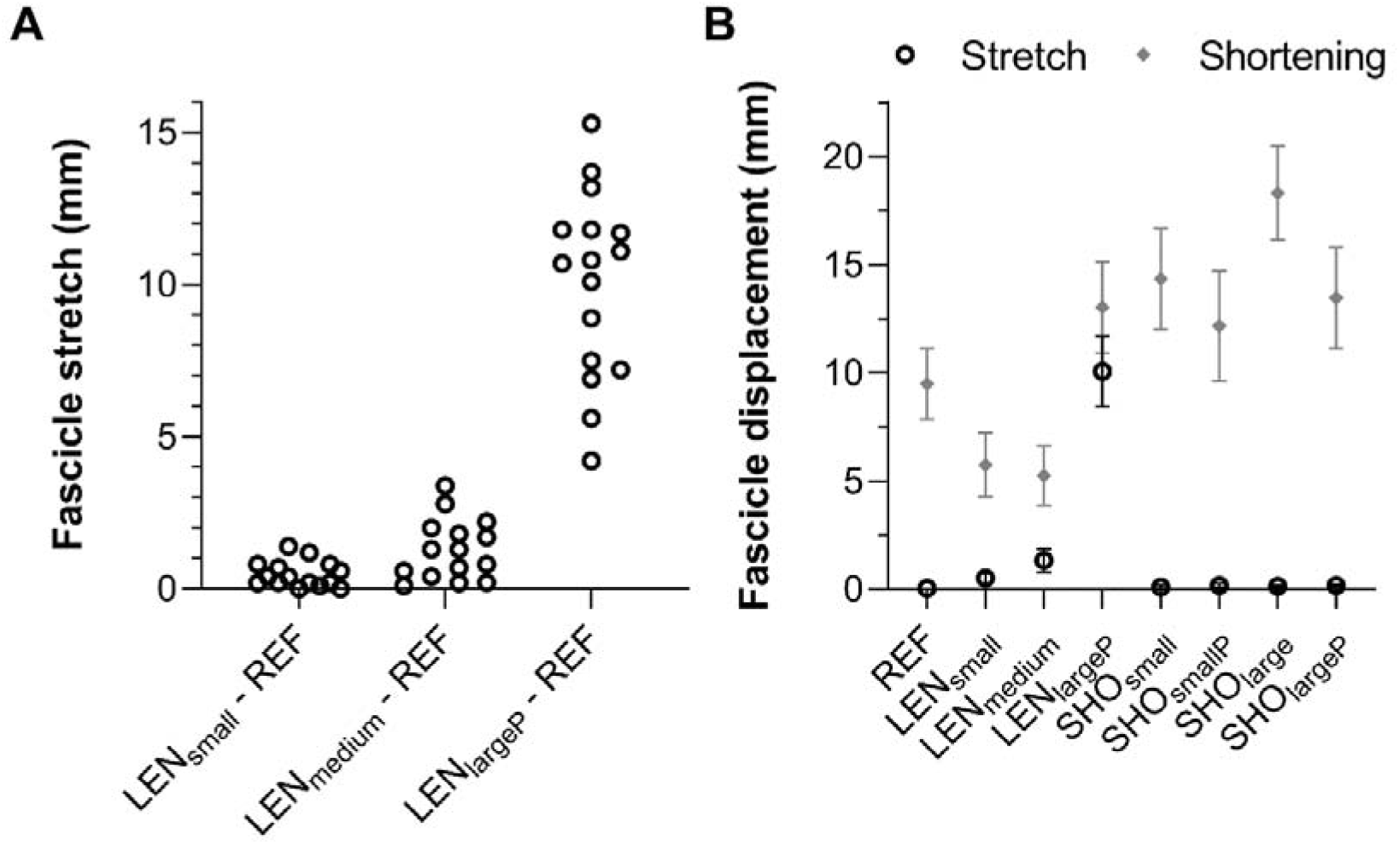
(A) Individual and (B) mean (± 95% confidence intervals) tibialis anterior fascicle (A&B) stretch (black open circles) and (B) shortening (grey filled diamonds) amplitudes during fixed-end reference (REF), and muscle-tendon unit lengthening (LEN)-hold and shortening (SHO)-hold conditions with and without a preload (_P_) over small, medium, and large amplitudes. The 95% confidence intervals show that fascicle shortening amplitudes were significantly different in the LEN and SHO conditions relative to REF.

The mean fascicle shortening amplitude was significantly different between conditions (*F*_2.72,40.06_ = 71.08, *p* < 0.001). The mean shortening amplitude was successfully reduced relative to REF in LEN_small_ (−3.7±1.5 mm, *p* < 0.001, *n* = 15) and LEN_medium_ (−4.2±1.9 mm, *p* < 0.001, *n* =15), but there was significantly more shortening relative to REF in LEN_largeP_ (3.6±2.0 mm, *p* < 0.001; Fig. 3B). As expected, there was also significantly more shortening relative to REF during MTU shortening (2.7-8.8 mm, *p* ≤ 0.001; Fig. 3B).

### Work production

Net muscle fascicle work production was significantly different between conditions (*F*_1.21,17.74_ = 65.47, *p* < 0.001). Mean net positive work was produced in LEN_small_ (0.3±0.5 J, *n* = 15), and this magnitude was significantly lower than in REF (1.4±0.6 J, *p* < 0.001; Fig. 4A). Mean net negative work was produced in LEN_medium_ (−0.4±0.7 J, *n* = 15) and LEN_largeP_ (−4.0±2.1 J), and more mean net positive work relative to REF was produced in SHO_small_ (2.4±1.4 J, *p* < 0.001), SHO_smallP_ (2.7±1.4 J, *p* < 0.001), SHO_large_ (3.3±1.6 J, *p* < 0.001), and SHO_largeP_ (3.6±1.5 J, *p* < 0.001; Fig. 4A). As different amounts of net negative fascicle work could potentially confound the relation between the steady-state active force percent differences (i.e. rFD) and net fascicle work production (Fig. 4A), and we were concerned with the relation between rFD and positive work, only the MTU shortening-hold conditions were considered in the following analysis. No significant repeated-measures linear relation was found between rFD and net fascicle work (*r*_rm_(47) = 0.16, 95% CI = −0.13 to 0.42, unadjusted *p* = 0.277).

**Figure 4.**
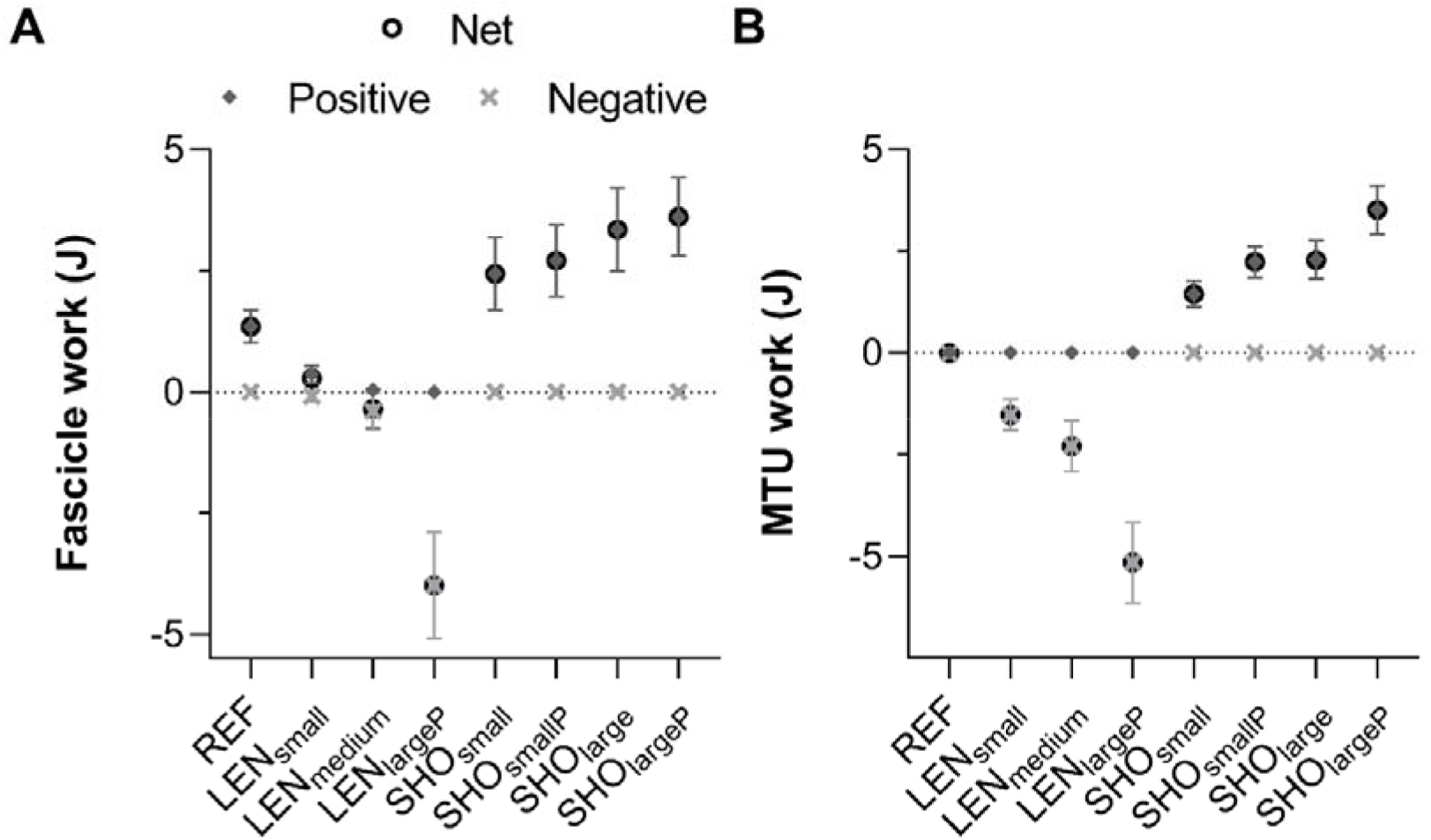
Mean (± 95% confidence intervals) tibialis anterior net (black open circles), positive (grey filled diamonds), and negative (light grey crosses) (A) fascicle and (B) muscle-tendon unit work magnitudes during fixed-end reference (REF), and muscle-tendon unit lengthening (LEN)-hold and shortening (SHO)-hold conditions with and without a preload (_P_) over small, medium, and large amplitudes. Active fascicle stretch in LEN_medium_ and LEN_largeP_ resulted in net negative fascicle work, but in LEN_small_ the relatively negligible active fascicle stretch resulted in net positive fascicle work. The 95% confidence intervals show that net fascicle and MTU work magnitudes were significantly different in the LEN and SHO conditions relative to REF.

Net MTU work production was significantly different between conditions (*F*_1.16,19.61_ = 131.4, *p* < 0.001). Zero net work was produced in REF (0.0±0.0 J; Fig. 4B). Net negative work was produced in LEN_small_ (−1.5±0.7 J, *n* = 15), LEN_medium_ (−2.3±1.1 J, *n* = 15), and LEN_largeP_ (−5.2±1.8 J), whereas net positive work was produced in SHO_small_ (1.4±0.6 J), SHO_smallP_ (2.2±0.7 J), SHO_large_ (2.3±0.9 J), and SHO_largeP_ (3.5±1.1 J; Fig. 4B). A significant repeated-measures linear relation was found between rFD and net MTU work (*r*_rm_(47) = 0.48, 95% CI = 0.24 to 0.67, unadjusted *p* < 0.001). Scatterplots of rFD relative to net fascicle and MTU work are shown in Figure 6.

**Figure 5.**
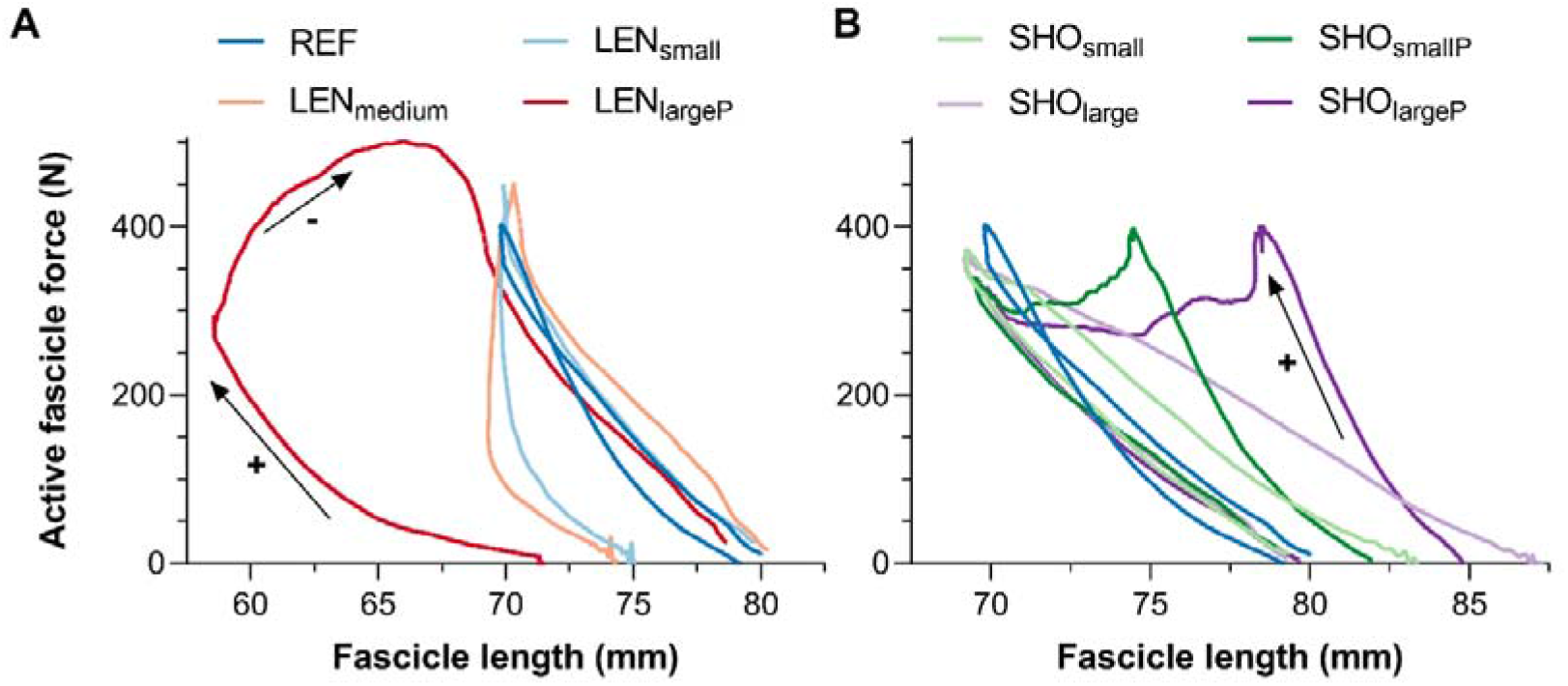
Mean tibialis anterior active fascicle force-fascicle length traces from sixteen participants during fixed-end reference (REF), and muscle-tendon unit (MTU) (A) lengthening (LEN)-hold and (B) shortening (SHO)-hold conditions with and without a preload (_P_) over small, medium, and large amplitudes in EMG-amplitude-matched contractions. The arrows indicate the directions of force production and fascicle displacement during the contractions, with shortening indicating positive work. Different active forces were clearly produced at similar fascicle lengths following MTU LEN and SHO. Also note the peak forces in SHO_smallP_ and SHO_largeP_ were attained prior to the imposed MTU SHO and there was 1 missing dataset for 2 separate participants in LEN_small_ and LEN_medium_.

**Figure 6.**
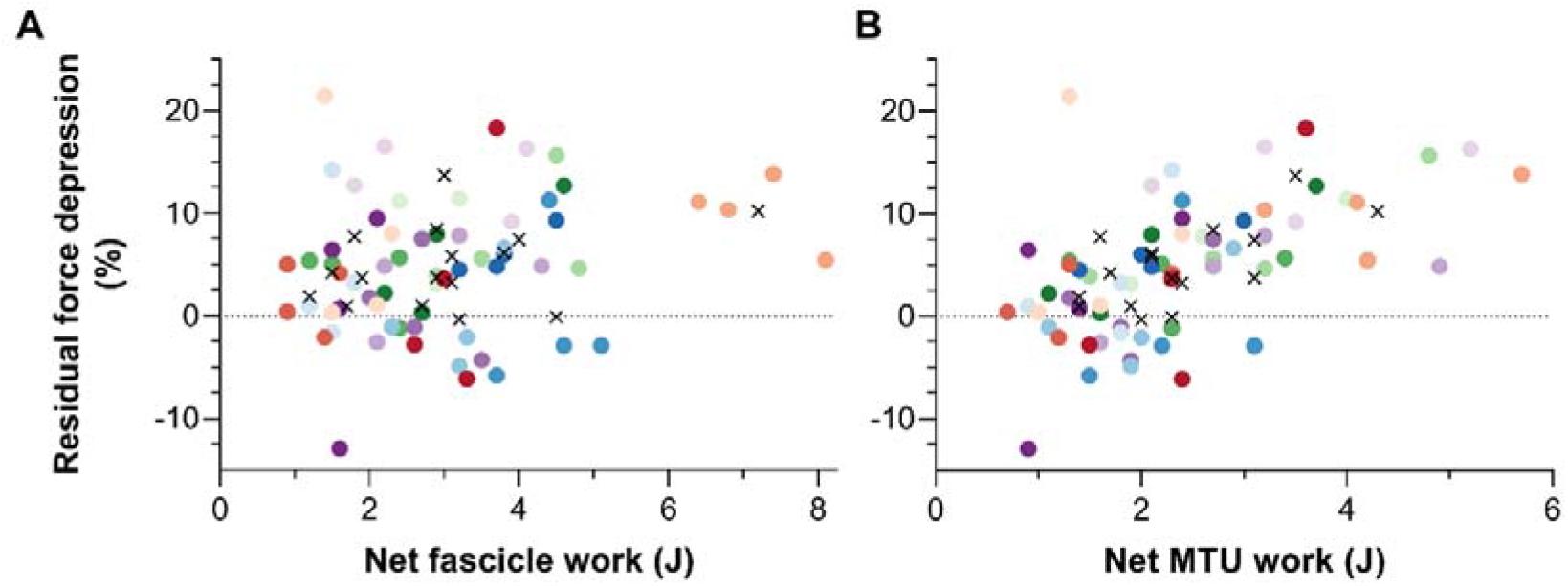
Scatterplots of tibialis anterior net (A) fascicle and (B) muscle-tendon unit work magnitudes relative to individual residual force depression magnitudes in the muscle-tendon unit (MTU) shortening (SHO)-hold conditions with and without a preload over small and large magnitudes. Each colour represents a different participant, and the black crosses represent the within-subject means across the four SHO conditions.

## Discussion

A muscle’s steady-state force following an active length change is difficult to predict because it is not solely dependent on the muscle’s activity level and degree of myofilament overlap. Instead, at given lengths and activity levels, a variable rFE or rFD can respectively exist following active lengthening or shortening (Abbott & Aubert, 1952). Active length changes therefore complicate the predictions of the subsequent muscle force. In this study, we compared steady-state active dorsiflexion torques and TA muscle fascicle force estimates following different amplitudes of MTU lengthening and shortening to the output attained during submaximal voluntary fixed-end dorsiflexion contractions at a matched MTU length and EMG amplitude. Despite similar steady-state TA muscle activity levels and muscle fascicle lengths between conditions, active torque and force were not similar in five out of seven conditions following the imposed MTU length change. Specifically, following MTU lengthening with and without a preload, active torque and force were enhanced by 3-5% relative to REF, but the enhancement was similar following small, medium, and large lengthening amplitudes. Following MTU shortening with a preload, active torque and force were depressed by 7-10% relative to REF, but the depression was not significantly different (1-2%) following small or large shortening amplitudes. Following MTU shortening without a preload, active torque and force were similar to REF following both small and large shortening amplitudes. These results support our first and second hypotheses that MTU lengthening enhances the subsequent active torque and force output while MTU shortening with a preload depresses it.

As predicted, we also found that rFD was strongly and positively correlated with MTU work during shortening, but not fascicle work. These findings indicate that at the relative muscle lengths we tested (i.e. approximately on the ascending limb of the active force-length relation, Raiteri *et al*., 2023), active dorsiflexion torque and TA fascicle force can be subsequently enhanced compared with REF regardless of whether the muscle fascicles are shortened or stretched during the imposed MTU lengthening. This level of enhancement could be exaggerated when compared with the MTU’s output during preloaded shortening because preloaded shortening increases MTU work during MTU shortening and depresses the subsequent MTU output. However, MTU shortening without a preload does not depress the subsequent MTU output more than a fixed-end contraction at a similar final MTU length and activity level.

### Muscle-tendon unit lengthening

A muscle that is actively lengthened produces a subsequently enhanced steady-state force relative to that produced during a fixed-end contraction at a similar final muscle length and activity level (Abbott & Aubert, 1952). In general, our findings align with previous *ex vivo* rFE findings from single sarcomeres (37-285%; Leonard *et al*., 2010), single myofibrils (106-386%; Joumaa *et al*., 2008), single muscle fibres (14±12%; Edman *et al*., 1982), and whole muscles (16±13%; Abbott & Aubert, 1952), as well as previous *in vivo* findings from the human dorsiflexors (9-16%; Tilp *et al*., 2009). However, we observed much less enhancement (3-5%; range: 0-15%), and this could be explained by the shorter relative final lengths we tested at, on the approximate ascending limb of TA’s active force-length relation (Raiteri *et al*., 2023). For example, Abbott & Aubert (1952) found rFE of only 7±7% (range: 0-16%) on the ascending limb and rFE of 27±17% (range: 13-52%) on the descending limb, and we previously showed that rFE increases with increasing muscle length (Bakenecker *et al*., 2022). Therefore, we expect that the enhancement we observed would have increased if we tested at longer relative final muscle lengths, if what we observed was indeed rFE.

One important difference between our study and many others that investigated whole muscles or intact MTUs is that we quantified fascicle kinematics of the prime mover during MTU lengthening. Had we not done this, we would have labelled the enhanced active torque and force output following all MTU lengthening conditions as rFE. However, as we observed net fascicle shortening during MTU lengthening without a preload, we believe that rFE-related mechanisms are unlikely to explain the enhanced steady-state output following small- and medium-amplitude MTU lengthening. This is because the mechanisms used to explain rFE, namely the reorganisation of a passive structural element (Herzog & Leonard, 2002; Herzog, 2018) and sarcomere length nonuniformity (Morgan, 1990; Edman, 2012), assume that all or some sarcomeres are stretched while activated. However, following small-amplitude MTU lengthening (i.e. LEN_small_), we observed net fascicle shortening (5.8±2.7 mm, range: 0.5-11.5 mm) and net positive fascicle work (0.3±0.5 J, range: −0.8 to 1 J), despite a small amount of active fascicle stretch (range: 0.1-1.4 mm) and negative fascicle work (−0.1±0.2 J, range: −0.8 to 0 J). Therefore, it is hard to envision how a passive structural element, which likely lies in parallel with the contractile elements (Edman *et al*., 1978, 1982; Herzog *et al*., 2003), could contribute enhanced forces when its length decreased during MTU lengthening. Additionally, as we ensured that MTU lengthening stopped at angles before the descending limb of the active force-angle relation (i.e. 15° plantar flexion, Raiteri *et al*., 2023), it is unlikely that sarcomere length nonuniformity largely contributed to an enhanced steady-state force on the ascending limb of TA’s active force-length relation. Furthermore, on the ascending limb, rFE decreases with increasing fascicle stretch amplitude (Hisey *et al*., 2009), but we did not observe a significant correlation between active steady-state force enhancement and fascicle stretch amplitude. For these reasons, we do not believe that the enhanced steady-state output we observed following MTU lengthening was due to rFE-related mechanisms.

We instead interpret the enhanced output following MTU lengthening with simultaneous net fascicle shortening or *lengthening* as ‘abolished’ rFD. This is because on the ascending limb of the active force-length relation, passive muscle force is typically not observed (Peterson *et al*., 2004; Herzog & Leonard, 2005), and despite increasing passive muscle force with increasing muscle length along with increasing passive force enhancement and rFE (Herzog & Leonard, 2005), rFE *decreases* with *increasing* stretch amplitude over the ascending limb (Hisey *et al*., 2009). This implies that a passive structural element engaged by cross-bridge cycling (Leonard & Herzog, 2010) does not contribute to the enhanced steady-state force following stretch over the ascending limb as rFE should either remain constant or increase with increasing stretch amplitude. Rather, we think that the fascicle shortening permitted due to in-series compliance during fixed-end contractions induces rFD, and that this rFD is abolished when the magnitude of fascicle shortening is reduced during MTU lengthening or when the muscle is actively stretched under negligible passive force. However, additional *ex vivo* experiments are needed to verify this interpretation, and to support it would ideally need to show a decrease in steady-state force, stiffness, and ATPase activity per second (Joumaa *et al*., 2017) in REF compared with following MTU lengthening.

The similar magnitudes of enhancement between LEN_small_ and LEN_medium_ suggests that additionally reducing fascicle shortening amplitude did not ‘reduce’ rFD more, and potentially implies that rFD is small (4.1±4.6%) within the human dorsiflexors during submaximal voluntary fixed-end contractions and MTU shortening-hold conditions without a preload. This level of enhancement relative to the REF condition following MTU lengthening with simultaneous net fascicle shortening agrees with what we previously observed (4%; Raiteri & Hahn, 2019), but this magnitude is roughly four times less than that reported by Mahmood and colleagues (18%; Mahmood *et al*., 2021). As only four rabbits were tested in that study, their reported enhancement is likely to be imprecise and may be an overestimate. Additionally, history-dependent effects might be larger during artificial stimulation than voluntary activation because during stimulation all muscle fibres are presumably activated (almost) simultaneously, whereas during voluntary activation muscle fibres are activated according to Henneman’s size principle (Henneman *et al*., 1965*a*, 1965*b*). Accordingly, higher threshold motor units presumably undergo less shortening as they are recruited later during active force development.

### Muscle-tendon unit shortening

A muscle that is actively shortened produces a subsequently depressed steady-state force relative to that produced during a fixed-end contraction at a similar final muscle length and activity level (Buchthal, 1942). Our findings align with previous *ex vivo* rFD findings from single sarcomeres (27±36%; Trecarten *et al*., 2015), single myofibrils (31±13%; Joumaa & Herzog, 2010), single muscle fibres (13±6%; Edman *et al*., 1993), and whole muscles (6&24%; Abbott & Aubert, 1952), as well as previous *in vivo* findings from the human dorsiflexors (9-20%; Tilp *et al*., 2009). We observed similar rFD following MTU shortening with a preload (7-10%; range: 0-21%), especially if we only consider rFD findings from the human dorsiflexors following slow-speed shortening (10°·s^-1^, 15° amplitude) under maximal voluntary activation (9±11%; Tilp *et al*., 2009). The larger rFD observed in *ex vivo* studies might be because the contractions investigated were artificially induced and activation was constant during shortening, which likely induced rFD in all muscle fibres during shortening, whereas under voluntary activation the level of activation might decrease (see Fig. 1, but bear in mind that activity level does not accurately reflect activation level), which could potentially eliminate rFD in some fibres. Additionally, if rFD exists in the fixed-end REF condition, this would act to reduce the steady-state active torque or force difference relative to any MTU shortening-hold condition. Indeed, once we substituted LEN_small_ as our reference condition to calculate rFD, which we did because we believe REF was contaminated with rFD and LEN_small_ was not, rFD of 13±6% was observed following large-amplitude MTU shortening with a preload, which is similar to the rFD observed in single frog muscle fibres following preloaded shortening under ∼75% of maximum tetanic force (13±6%; Edman *et al*., 1993). Similar rFD therefore appears to exist across muscle scales following MTU shortening with a preload.

Similar rFD also appears to exist across muscle scales following MTU shortening without a preload. We observed similar steady-state active torque and force output in our fixed-end REF, SHO_small_, and SHO_large_ conditions, which is in line with observations from frog muscle fibres following isotonic shortening against a relatively light (≤30% of maximum) load (Edman, 1966; Gordon *et al*., 1966). This is interesting because *ex vivo*, all frog fibres were presumably activated during light-load isotonic shortening before isometric force development (Edman, 1966; Gordon *et al*., 1966), whereas *in vivo*, new fibres were likely recruited during shortening according to Henneman’s size principle (Henneman *et al*., 1965*a*, 1965*b*); Consequently, the activation pattern during the initial shortening accompanying active force development does not seem to have a measurable effect on rFD.

Edman (1966) and Gordon and colleagues (1966) suggested that previous isotonic shortening does not impact the muscle’s capacity to produce force isometrically with a given amount of myofilament overlap. However, this interpretation was based on the assumption that the steady-state reference force was not depressed. We think our findings suggest otherwise; the fixed-end REF contractions and MTU shortening-hold contractions without a preload were depressed by 3-5% relative to conditions with less fascicle shortening. Hill (1949) also observed that muscle force could be elevated by either applying a quick stretch early following a single maximal stimulation to reduce shortening, or by applying a successive maximal stimulation once the in-series elastic tissues were stretched by the first stimulation. Additionally, Mutungi and Ranatunga (2000) found two to three times higher peak twitch force outputs from slow and fast rat muscle fibre bundles, respectively, by reducing the extent of sarcomere shortening by 60% or more, and found lower enhancement at longer sarcomere lengths with less sarcomere shortening during fixed-end contractions. Therefore, in line with a later suggestion by Edman (1975) that active sarcomere shortening of a critical amplitude can be expected to measurably reduce steady-state force, we would like to emphasise the importance of considering shortening-induced rFD during voluntary fixed-end contractions.

### Mechanisms

Although *in vivo* observations cannot shed light on the molecular mechanisms underlying rFD, we think that it is worth speculating about these mechanisms to help formulate predictions across muscle scales. Hill (1951) explained that the depression in peak force during a fixed-end twitch following the addition of series compliance was due to the wasted time spent stretching the series elastic element. Later, Edman (1975) suggested that a peak twitch force depression following shortening was compatible with a structural change of the myofilament system during active shortening. More specifically, he proposed that the physical state of the troponin-tropomyosin complex along the thin filament was possibly affected during shortening, leading to a transitory impairment of actin-myosin interaction. Although later research showed the skeletal troponin C structure is unlikely to be affected by cycling cross bridges in the presence of Ca^2+^ (Martyn *et al*., 1999), this mechanism is similar to the perhaps most-commonly cited mechanism underlying rFD, which is a stress-induced inhibition of cross bridges entering the new overlap region during shortening (Maréchal & Plaghki, 1979).

This stress-induced cross-bridge inhibition mechanism was later formalised (Herzog, 1998) and built upon to also include a stress-induced inhibition of cross bridges in the old overlap region before the imposed shortening (Joumaa *et al*., 2012). This addition was made because if only new cross bridges entering the overlap zone were affected during shortening, rFD should not exceed the force difference between the initial length on the descending limb of the active force-length relation and the final length, which is what was originally found (Maréchal & Plaghki, 1979). However, later research showed that rFD can exceed the force difference between the initial length on the descending limb and the final length (Granzier & Pollack, 1989; Joumaa & Herzog, 2010; Joumaa *et al*., 2012). We discuss the predictive ability of this updated theory below.

One prediction of the stress-induced cross-bridge inhibition theory is that rFD increases with increasing shortening amplitude due to a larger new overlap region. While this prediction is generally supported by experimental findings (Abbott & Aubert, 1952; Maréchal & Plaghki, 1979; Sugi & Tsuchiya, 1988; Granzier & Pollack, 1989; Herzog & Leonard, 1997; De Ruiter *et al*., 1998; Morgan *et al*., 2000; Lee & Herzog, 2003; Schachar *et al*., 2004; Bullimore *et al*., 2007), other predictions have mixed experimental support. For example, steady-state force reductions were similar following shortening at various times during the rising phase of the twitch (Edman, 1975), and rFD was similar after changing mid-shortening from a slow (2.5 mm·s^-1^) to fast (25.6 mm·s^-1^) or fast to slow speed (Maréchal & Plaghki, 1979). However, the stress-induced cross-bridge inhibition theory predicts that higher forces during constant-amplitude shortening should induce greater rFD because of higher stress on actin. Additionally, rFD remained when fast-speed (200 mm·s^-1^) shortening, which caused force to drop to zero, was preceded by slow-speed (4.5 mm·s^-1^) shortening (Herzog & Leonard, 2007), despite the theory predicting that releasing the stress on actin should abolish rFD. Consequently, rFD cannot be solely explained by the stress-induced inhibition theory.

Similarly, rFD cannot be solely explained by non-uniform sarcomere lengths following shortening because rFD has been observed on the ascending limb of the active force-length relation (Herzog *et al*., 1998; Pun *et al*., 2010; McDaniel *et al*., 2010*a*), and experiments on single myofibrils (Joumaa and Herzog, 2010; Pun et al., 2010) and single sarcomeres (Trecarten *et al*., 2015) showed that rFD can occur without significant increases in non-uniformity relative to the fixed-end REF condition. However, more pronounced sarcomere length non-uniformity might partly explain the greater rFD following slow-speed compared with fast-speed shortening (Sugi & Tsuchiya, 1988; Edman *et al*., 1993). A role of passive elements, such as titin, could also contribute to rFD by interacting with the thin filament during activation (Kellermayer & Granzier, 1996; Nagy *et al*., 2004) and interfering with myosin-binding sites during and following shortening (Rode *et al*., 2009; Nishikawa *et al*., 2012). Unloading of titin and lower titin-based passive forces during shortening might also limit thick- and thin-filament activation following shortening (Cazorla *et al*., 2001). However, at present, titin-based contributions to rFD have limited experimental support (Tahir *et al*., 2020) and it is unclear, at least to us, how titin-based mechanisms could contribute to rFD when shortening occurs exclusively on the ascending limb of the active force-length relation, where passive forces are minimal (McDaniel *et al*., 2010*a*). Perhaps other parallel elements in the A-band, such as myosin-binding protein C (Offer, 1973), bind to actin (Luther *et al*., 2011) and induce rFD by interfering with filament sliding (Robinett *et al*., 2019) at relatively short sarcomere lengths, but this mechanism requires additional experimental support before it should be more seriously considered.

Based on the available data, we think that the most plausible mechanism underlying rFD is an increase in cross-bridge detachment rate during shortening that reduces the number of attached cross bridges and fraction in a strongly-bound state following shortening. Earlier cross-bridge detachment during shortening (Huxley *et al*., 2006) would decrease the cross-bridge duty cycle (i.e. time attached relative to the cycle duration), and reduce the number of attached cross bridges (Huxley, 1957) and fraction in a strongly-bound state, decreasing force and increasing the ATP consumption rate relative to isometric conditions (He *et al*., 2000). We speculate that less time spent shortening, either because the imposed shortening amplitude is small or the speed is fast, reduces the number of cross bridges that contribute to shortening (Linari *et al*., 2015; Fusi *et al*., 2017), and the subsequent cross-bridge inhibition following shortening. This theory could explain why rFD is not measurable following fast-speed (200 mm·s^-1^) shortening (Herzog & Leonard, 2007) because supposedly only a very low fraction (5%) of cross bridges contribute to unloaded shortening (Linari *et al*., 2015; Fusi *et al*., 2017), which might not heavily affect net cross-bridge cycling following shortening and force redevelopment.

A faster cross-bridge detachment rate during shortening has some support from multiscale 3D lattice model findings, which suggest that a large number of cross bridges initially rapidly transition from a weakly-bound to a strongly-bound state and then detach quickly (Mijailovich *et al*., 2016). Subsequently, once a steady-state shortening speed is achieved, the net number of attached cross bridges and fraction in a strongly-bound state is reduced (Mijailovich *et al*., 2016). These changes could persist following shortening because of reduced cooperativity among fewer strongly-bound cross bridges, and could subsequently lead to reduced force per cross bridge and reduced thin filament activation (Caremani *et al*., 2022; Brunello *et al*., 2023). Notably, these changes have some indirect experimental support (Joumaa *et al*., 2021), and potentially explain the lower rFD during activation via MgADP versus Ca^2+^ (Pun *et al*., 2010; Trecarten *et al*., 2015), because MgADP biases cross bridges into a strongly-bound state (Kintses *et al*., 2007; Awinda *et al*., 2022). However, additional spatially-explicit modelling and small-angle X-ray diffraction data are needed to gain further insight about this mechanism, particularly regarding what happens to cross-bridge detachment rates and the resulting cross-bridge state distributions when shortening amplitude and speed are systematically manipulated.

The final potential mechanism underlying rFD that we would like to discuss is a work-induced rather than stress-induced inhibition of cross-bridge attachment (Maréchal & Plaghki, 1979; Herzog *et al*., 2000). This hypothesis explains observations that rFD is positively and linearly related to the shortening amplitude (Maréchal & Plaghki, 1979; Schachar *et al*., 2004; Bullimore *et al*., 2007), as well as the mean force during shortening (Granzier & Pollack, 1989; De Ruiter *et al*., 1998; Herzog *et al*., 1998). However, the rFD-work relation is not necessarily linear (Herzog *et al*., 2000), only positive MTU work rather than positive muscle work seems to matter based on our results, and similar or different rFD can be achieved following vastly different or similar amounts of positive muscle work, respectively (Leonard & Herzog, 2005; Van Noten & Van Leemputte, 2011). Additionally, despite increased positive muscle work, rFD is not larger following a stretch-shortening cycle than following preloaded MTU shortening (Seiberl *et al*., 2015; Fortuna *et al*., 2017, 2018, 2019; Hahn & Riedel, 2018; Tomalka *et al*., 2020; Groeber *et al*., 2020, 2021; Fukutani & Herzog, 2021), nor is rFD larger following a two-step than one-step shortening contraction (Herzog & Leonard, 2000). These previous results in combination with our findings suggest that rFD is only linearly related to positive muscle work under some specific conditions. These conditions include shortening over different amplitudes, but at a constant speed, and shortening over the same amplitude, but at different speeds. Clearly, these conditions are unlikely to occur during everyday movement. Additionally, predicting rFD based on MTU work would lead to inaccuracies following MTU shortening without a preload because there would be substantial net positive work production, but no increased rFD relative to a fixed-end condition with net zero work production. Another factor besides work is thus needed to accurately predict rFD.

We speculate that rFD is strongly and linearly related to the number of active muscle fibres during shortening. Although we cannot estimate motor unit recruitment from the bipolar surface EMG measurements used in our experiments, we explored the repeated-measures linear relation between rFD and the mean muscle activity level during shortening between the MTU shortening-hold conditions and found it was significant (*r*_rm_(47) = 0.63, 95% CI [0.43 to 0.78], unadjusted *p* < 0.001). As the muscle activity level during shortening was similar between REF and the MTU shortening-hold conditions without a preload, this variable helps to explain why rFD was similar between these conditions. Additionally, as the muscle activity level was systematically higher during preloaded MTU shortening, but similar between preloaded shortening-hold conditions of small and large amplitudes, the mean muscle activity level during shortening also helps to explain why rFD was not significantly affected by shortening amplitude. Fewer active fibres during shortening should thus reduce rFD following shortening, which has been previously reported following reductions in stimulation frequency or stimulation intensity during shortening (Herzog & Leonard, 1997; De Ruiter *et al*., 1998; Herzog *et al*., 1998). However, increasing the number of active fibres during shortening compared with the steady state should not necessarily increase rFD because the additionally-recruited fibres should lose their rFD when they are derecruited following shortening. Further, the mean force during constant-amplitude shortening is potentially linearly related to rFD because during slow-speed shortening, additional fibres (i.e. low- and high-threshold motor units) have cross bridges that can contribute to shortening and are subsequently inhibited following shortening, whereas during fast-speed shortening, only fibres with high enough myosin ATPase activity (i.e. high-threshold motor units only) can contribute to shortening (Bárány, 1967) and are subsequently inhibited. Nevertheless, despite the twenty-five-years-old suggestion that rFD is linearly related to the number of active fibres during shortening (Herzog *et al*., 1998), future experiments are still needed to verify the validity of this hypothesis.

### Implications

rFD is often considered as a negative consequence of muscle shortening because it depresses contractile output. However, here we show that the rFD during fixed-end contractions and MTU shortening without a preload is small (3-5%), and that rFD can be reduced and potentially abolished by timing MTU lengthening with active force development. Interestingly, the timing and rate of MTU lengthening coincide with the onset of muscle activity and its rate of increase in relatively compliant MTUs during efficient everyday tasks such as walking and running (Lichtwark & Wilson, 2006). A tight coupling between the timing of MTU lengthening and muscle activation might be important for minimising energy expenditure in relatively compliant MTUs because MTU lengthening reduces the muscle’s internal shortening requirement by effectively stiffening the series elastic element, which subsequently increases the force output for a given muscle activity level. These mechanics might also be more favourable than stretch-shortening mechanics at the muscle fascicle level because as long as similar extra forces are attained during MTU lengthening, this additional energy could be stored by the stretched series elastic element and later reused during MTU shortening, instead of being lost as heat during active muscle stretch.

The increased rFD observed following MTU shortening with a preload might not be that physiologically relevant because everyday movements are usually not, if ever, preceded by a fixed-end preload. However, the mechanism underlying the observation could be physiologically important, especially for muscle relaxation. This is because the rate of force relaxation is generally slower than the rate of force development (Rome *et al*., 2000; Tesi *et al*., 2002), which limits positive muscle work output during the brief, cyclic contractions that typically power locomotion (McDaniel *et al*., 2010*b*). But importantly, force relaxation can be accelerated when shortening occurs simultaneously, which reduces energy expenditure (Lou *et al*., 1998) and amplifies the series elastic element’s power output (Barclay & Lichtwark, 2007). As faster shortening velocities further accelerate the rate of relaxation (Barclay & Lichtwark, 2007), faster stride frequencies are potentially achievable during locomotion due to an earlier return to a resting muscle state. This idea is consistent with findings from R146N mutant myosin, which produced less work and power during cycling contractions because of a slower ATP-induced detachment rate from actin and thus a longer-duration strongly-bound cross-bridge state (Kronert *et al*., 2018). Consequently, muscle shortening amplifies MTU power output and reduces energy expenditure during muscle relaxation, and the mechanism or mechanisms underlying this shortening-speed-dependent relaxation rate might be similar to those underlying rFD.

The higher steady-state force output we observed following MTU lengthening, and the lower steady-state force output we observed following preloaded MTU shortening, add to the growing body of evidence that inaccurate force predictions from muscle models are likely to occur when contraction history is neglected. However, based on the mean enhancement and depression values of 3-5% and 7-10% we respectively observed, it could be argued that accounting for contraction history is not worthwhile because force prediction errors of *at least* 10% of maximum active isometric force have been reported from muscle models during movement simulations (Lee *et al*., 2013; Dick *et al*., 2017). To us, this argument is shortsighted, because contraction history affects forces more during than following movement (30% versus 5%; McDaniel *et al*., 2010*a*). Consequently, future research should investigate how forces during lengthening and shortening are affected by contraction history. We hope that our experimental approach and data help to respectively guide and motivate studies on this topic, as well as cautions modellers against using muscle work to predict forces during and following active shortening.

### Limitations

Although we observed similar steady-state TA muscle activity levels between our eight conditions, this does not necessarily indicate there were similar levels of neural drive to the TA in each condition (Farina *et al*., 2004). However, as we performed repeated measurements within the same session and the desired activity levels were submaximal (∼25% of the angle-specific maximum), the degree of amplitude cancellation was likely to be small (Keenan *et al*., 2005). Therefore, we believe it is reasonable to assume that TA activation was similar at matched MTU lengths when the steady-state torque and force outputs were quantified. We also believe that synergistic activation was likely to be similar between our eight conditions because of the similar steady-state TA fascicle lengths we observed. While it could be argued that a major limitation was that we did not measure and correct for plantar flexor muscle co-contraction, we have data from two different ultrasound imaging techniques that suggest plantar flexor contributions to the measured steady-state net ankle joint torque are likely to be negligible and that increases in plantar flexor surface EMG amplitudes during fixed-end dorsiflexion contractions are primarily due to increasing cross-talk from the dorsiflexors (Raiteri *et al*., 2015, 2016). Consequently, these previous studies give us confidence that the active dorsiflexion torque differences we observed between conditions were not due to plantar flexor muscle co-contraction differences.

We were unable to completely eliminate active fascicle stretch in all participants in the MTU lengthening-hold conditions without a preload and it could therefore be argued that LEN_small_ and LEN_medium_ induced rFE. However, we think this is unlikely because the amplitudes of fascicle stretch did not exceed 0.4 mm in 9 out of 16 participants or 0.1 mm in 3 out of 16 participants, yet the steady-state active forces were still enhanced by 5±4% and 4±5%, respectively, which was similar to the 3±4% enhancement in the remaining participants who showed fascicle stretch magnitudes ranging from 0.6-3.4 mm. Indeed, the probability of actively stretching sarcomeres was likely increased during MTU lengthening compared with the other conditions, which could have increased the probability of observing rFE following MTU lengthening, but then we would have expected a significant and positive linear repeated-measures relation between the enhanced steady-state force output and fascicle stretch amplitude, and this was not the case. Similarly, increased fascicle stretch amplitudes within the two-dimensional area of the TA that was imaged likely increased the probability of stretch in other non-imaged areas of the TA, as well as the other dorsiflexors, but this clearly did not result in greater enhancement following MTU lengthening, which suggests to us that negligible rFE was induced following MTU lengthening. However, steady-state stiffness measurements from isolated muscle preparations are needed to quantify whether rFE (i.e. similar/increased stiffness relative to REF) or rFD (i.e. decreased stiffness) exists following MTU lengthening.

## Conclusions

This study was designed to determine whether shortening during the initial force rise of fixed-end dorsiflexion contractions (i.e. REF) induces rFD and to test whether the relation between rFD and positive muscle or MTU work is strong and linear under submaximal voluntary activation in healthy women and men. MTU lengthening-hold conditions over increasing amplitudes and with a preload reduced the amount of active TA fascicle shortening during MTU lengthening by increasing active fascicle stretch amplitudes, but the magnitudes of active fascicle force enhancement following MTU lengthening relative to REF were similar at 3-5%. In particular, the steady-state force enhancement of 3±5% following small-amplitude MTU lengthening without a preload suggests that REF was contaminated with rFD because of greater shortening-induced force loss during force development due to effectively greater in-series compliance. MTU shortening-hold conditions over increasing amplitudes and with a preload increased both the amount of active TA fascicle shortening and positive fascicle work during the contractions, but the magnitudes of active fascicle force depression following MTU shortening relative to REF could not be accurately predicted based on fascicle work. While MTU work provided more accurate predictions of rFD, the within-subjects linear relation was only moderate and MTU work did not predict similar rFD magnitudes between REF and the MTU shortening-hold conditions without a preload. We speculate that the number of muscle fibres active during shortening is strongly correlated with rFD, as well as dynamic force depression during shortening. We hope that our study motivates others to investigate how contraction history affects neuromuscular output, which is sorely needed to help validate and inform the muscle models used in everyday movement simulations.

## Data availability statement

The data from this study will be made available before publication.

## Author contributions

B.J.R and D.H contributed to conception and design of the study and interpretation of data; B.J.R and L.L contributed to acquisition and analysis of data; B.J.R drafted the manuscript; all authors contributed to revising the manuscript critically for important intellectual content and approved the final version of the manuscript. All authors agree to be held accountable for all aspects of the work, those designated as authors qualify for authorship, and those who qualify for authorship are listed.

